# Anthropogenic noise affects vocalisation properties of the territorial song of Western Australian magpies

**DOI:** 10.1101/2024.02.15.578887

**Authors:** Grace Blackburn, Mylene Dutour, Benjamin J. Ashton, Alex Thornton, Amanda R. Ridley

## Abstract

Anthropogenic noise is considered one of the most serious forms of pollution globally and has been shown to have negative effects on the distribution, behaviour, cognition and reproductive success of animal species worldwide. One of the most well researched impacts of anthropogenic noise is its effect on acoustic communication. Animals may adjust the rate, amplitude, duration, and/or frequency of their acoustic signals to better maintain communication when anthropogenic noise is present. In this study, we combine behavioural focals, amplitude measurements and audio recordings to investigate how female Western Australian magpies (*Gymnorhina tibicen dorsalis*) alter the acoustic features of their territorial song (known as a carol) when anthropogenic noise is present. Magpies reduced the rate at which they carolled when loud anthropogenic noise was present, and increased their peak frequency during anthropogenic noise, but, contrary to our hypotheses did not alter the amplitude, duration, or other frequency parameters of their carols. Results from this study add to the growing body of literature documenting changes to the vocal behaviour of wildlife in the presence of anthropogenic noise and highlights the importance of investigating multiple acoustic parameters to understand how species adjust their vocalisations in response to this pollutant.

## Introduction

Recent urbanisation and human expansion into previously untouched areas has led to a significant increase in the amount of noise generated by human activities (Barber et al., 2010; Shannon et al., 2016). Such increases in anthropogenic noise have been shown to have a range of negative effects on both human and other species (Kunc & Schmidt, 2019; Shannon et al., 2016; World Health Organization. Regional Office for Europe, 2011). In humans, sleep deprivation, cardiovascular disease, hearing loss and cognitive impairment have all been linked to anthropogenic noise (World Health Organization. Regional Office for Europe, 2011). In other animals, declines in species abundance and richness (Goodwin & Shriver, 2011; McClure et al., 2013; Wilson et al., 2021), as well as changes in behaviour (Blackburn et al., 2023; Eastcott et al., 2020; Evans et al., 2018), cognition (Osbrink et al., 2021; Templeton et al., 2023), communication (Duquette et al., 2021; Kunc & Schmidt, 2021), and reductions in reproductive success (Mulholland et al., 2018; Schroeder et al., 2012) have been linked to anthropogenic noise.

Research into the effects of anthropogenic noise on wildlife has largely focused on the effects of noise on acoustic communication, or rather, the ways in which animals modify their acoustic signals in response to anthropogenic noise (Duquette et al., 2021; Kunc & Schmidt, 2019; Shannon et al., 2016). Noise-related disruptions or alterations to acoustic signals have been documented in many species and environments, with four common changes typically observed. First, animals may alter the rate of their calling such that they vocalise less or avoid vocalising all together when noise is present, and thus minimise the energetic costs associated with vocalising when these vocalisations may be partially or completely masked by anthropogenic noise (Hart et al., 2015; Pearson & Clarke, 2018). Deemed the ‘silentium effect’ by Pearson and Clarke (2018), declines in vocalisation rate due to anthropogenic noise have been reported in anurans, whales, bats, and birds (Blackburn et al., 2023; Gross et al., 2010; Lengagne, 2008; Melcón et al., 2012; Pearson & Clarke, 2018; Tsujii et al., 2018). A second and much more commonly reported alteration to animals’ acoustic signals is the Lombard effect (Brumm & Zollinger, 2011; Kunc et al., 2022), where individuals increase the amplitude of their vocalisations so as to maintain broadcast distance in noisy conditions. Initially described in humans (Lombard, 1911), the Lombard effect is now considered a widespread phenomenon among animal species (Brumm & Zollinger, 2011). Third, animals may change the frequency of their vocalisations, generally by increasing the frequency of their calls to avoid or minimise acoustic masking by low frequency anthropogenic noise (Nemeth & Brumm, 2009; Proppe et al., 2011; Roca et al., 2016). Recent meta-analyses have found that terrestrial species generally increase the minimum but not the maximum or peak frequencies of their vocalisations, as it is the lower frequency elements of vocalisations that are most likely to be affected by anthropogenic noise (Duquette et al., 2021; Kunc & Schmidt, 2021). Finally, individuals may alter the duration of their vocalisations during anthropogenic noise (Francis et al., 2011; Gomes & Goerlitz, 2020; Kunc & Schmidt, 2021; Potvin & Mulder, 2013), generally by increasing the length of their vocalisation to improve signal detection in the presence of increased noise (Brumm et al., 2004; Francis et al., 2011). Despite changes to the rate, amplitude, frequency, and duration of vocalisations being attributed to anthropogenic noise, few studies consider *all* of these parameters simultaneously (Nemeth & Brumm, 2009; Pearson & Clarke, 2018; Proppe et al., 2011; Sementili–Cardoso & Donatelli, 2021; Slabbekoorn & den Boer-Visser, 2006; Templeton et al., 2016).

Here, we use behavioural focals, amplitude measurements, and audio recordings to investigate how female Western Australian magpies vary the acoustic parameters of their carols when anthropogenic noise is present. Western Australian magpies are a highly vocal subspecies of Australian magpie found in urban areas in the south-west of Western Australia (Dutour & Ridley, 2020; Johnstone, 2004). Magpies are highly territorial, and produce multiple complex vocalisations (Dutour & Ridley, 2020; Dutour et al., 2020; Walsh et al., 2023). The main territorial song of magpies – known as “carols” – are high-amplitude, slurred songs sung by individuals to advertise territory ownership and defend territories (Blackburn et al., 2022; Carrick, 1963; Dutour & Ridley, 2020). Previous work on this sub-species revealed that females carol more often than males (Dutour & Ridley, 2020), therefore in this study we focused on female carols. We predict that magpies will decrease the rate at which they carol, and increase the amplitude, duration, and minimum frequency of their carols when anthropogenic noise is present, to maintain their communication in the face of anthropogenic noise.

## Methods

### Study species and site

Western Australian magpies live in cooperative breeding groups ranging in size from 2 to 12 individuals and use vocal cues to communicate with group members about predators and other threats, defend territories, and maintain group cohesion (Dutour & Ridley, 2020; Dutour et al., 2020; Pike et al., 2019). Individuals in this subspecies are sexually dichromatic, allowing sex to be visually discerned once individuals achieve their adult plumage, at roughly three years of age (Dutour & Ridley, 2020). Western Australian magpies are well adapted to urban areas and often reside in urban suburbs and parklands around Perth, and hence are exposed to anthropogenic noise daily, from sources such as road traffic, planes, trains, construction, and/or machinery (Blackburn et al., 2023; Johnstone, 2004).

Data were collected from July 2021 to August 2023, between 5-11am (when birds are most active (Edwards et al., 2015)) on 15 groups of magpies (group sizes ranged from 2 – 10 adults during this study) located in urban areas of Crawley and Guildford, Western Australia. Birds in this population are habituated to human presence, allowing close observation and recording of natural behaviour, and individuals are identifiable via coloured rings, plumage anomalies, or distinctive scarring (Ashton et al., 2022).

### Behavioural focals

To assess the effect of anthropogenic noise on the carol rate of magpies, we conducted 20-minute focal observations on female magpies between October 2021 and February 2022 (*N* = 126 focals across 39 birds from 15 groups; mean ± SE = 3.23 ± 0.24 focals per bird) noting all behaviours and vocalisations (including carols) that individuals produced. During focals, the presence of anthropogenic noise was recorded, and a DIGITECH Micro Sound Pressure Level Meter (hereafter, SPL meter) (frequency weighting: A, time weighting: F) was used to measure the maximum amplitude of these noise events. Noise conditions during the focals were classified as either ‘loud anthropogenic noise’ (LAN), if the maximum amplitude exceeded 50dB, or ‘background noise’ (BN), if the maximum amplitude did not exceed 50dB. This threshold of 50dB has been used in previous studies to differentiate levels of anthropogenic noise (Blackburn et al., 2023; Grade & Sieving, 2016). Each focal was split into time when noise was classified as ‘loud anthropogenic noise’ and time when noise was classified as ‘background noise’, allowing for calculation and comparison of the carol rates of magpies during both noise conditions. A customised ethogram was created in CyberTracker (CyberTracker Conservation 2021) to record the behaviour of individuals to the nearest second.

### Amplitude measurements

To determine the effect of anthropogenic noise on the amplitude at which female magpies carol, we used a SPL meter to record the maximum sound pressure level (SPL) of naturally emitted carols. For each carol, we noted the identity of the caller, the group the individual was in, distance from the bird to the sound pressure level meter (m), orientation of the bird (whether it was facing towards, side on or away from the sound pressure level meter), location of the bird (whether it was above, below, or in line with the sound pressure level meter), the number of conspecific individuals present within 15 meters (because social context and proximity is thought to affect the amplitude at which individuals vocalise (Brumm & Ritschard, 2011; Brumm & Zollinger, 2011)), the weather condition (cloudy or clear), and anthropogenic noise condition (LAN if background noise was > 50dB, and BN if background noise was < 50dB). Humidity and temperature are also thought to affect signal transmission and attenuation (with higher temperatures and lower humidity generally resulting in greater signal attenuation, though these effects are dependent on other attributes of the signal and environmental conditions (Brumm & Zollinger, 2011; Goerlitz, 2018; Wiley & Richards, 1978)) and therefore may affect measurements of amplitude. Therefore, temperature and humidity data were retrieved from the Bureau of Meteorology records from the nearest weather station, ∼4km away from the Guildford field site, and ∼5km away from the Crawley field site (Australian Government Bureau of Meteorology, 2023) and included in the analysis. Only measurements taken from adult females, at a distance of 8m or less, and with birds orientated towards and level with or above the SPL meter were included in the analysis (*N* = 71 amplitude measurements, from 17 birds from 10 groups). Amplitude measurements were only taken if the path between the SPL meter and the bird was clear of any tree branches or other obstacles. For each carol emitted by the focal bird, we collected a measurement of the maximum SPL of the carol, and a measurement of the SPL directly after the carol, to use as a measure of the background noise SPL occurring during the carol. We then followed the logarithmic computational formula detailed in (Brumm & Zollinger, 2011) to subtract the background noise SPL from the carol SPL, in order to gain a measure of the SPL of the carol itself. We then used the logarithmic formula detailed in Brumm and Zollinger (2011) to standardise the amplitude of carols to a distance of one meter.

### Song recordings

To determine the effect of anthropogenic noise on the duration and frequency of female carols, we recorded naturally produced carols using a RODE NTG-2 directional microphone set within a Blimp suspension windshield system and attached to a Roland R-07 wave/MP3 recorder at a sampling rate of 44.1kHz. For each carol, individual and group identity were recorded, as well as the distance between the bird and microphone, and the presence or absence of any anthropogenic noise event (presence and absence of discernible anthropogenic noise was noted by the experimenter during each carol) as well as the type of anthropogenic noise that was occurring during the carol (traffic noise, plane noise, machinery and train noise). Only carols that did not overlap with the songs of other birds and were recorded from a distance of less than 8 meters were included in the analysis. Females were only included in the analysis if we had recordings of their carols both when discernible anthropogenic noise events were present *and* when they were absent.

### Acoustic analysis

Avisoft SASLab (version 5.3.2) was used to measure duration and frequency parameters of 64 carols from 15 female magpies across 11 groups. The duration of each carol was measured manually from spectrograms. Prior to frequency analysis, we conducted high-pass filtering of recordings (cut-off frequency: 0.5 kHz) to increase signal-to-noise ratio (Templeton et al., 2016). We then created an amplitude spectrum (linear) for each carol (Hamming window, using the function ‘Normalize to maximum’) and used the automated spectral characteristics function to determine the following acoustic features from carols: (1) peak frequency; (2) maximum frequency; (3) minimum frequency; and (4) frequency bandwidth. A threshold of -5dB (chosen as all carol peak frequencies were at least 5dB above the background noise in our recordings) below the peak frequency was used to measure frequency parameters.

### Statistical analysis

The effect of anthropogenic noise on magpie carolling behaviour and acoustic features was analysed via model selection using linear mixed models (LMMs) and generalized linear mixed models (GLMMs) in the *glmmTMB* package (Brooks et al., 2017) in R studio 4.2.0 (RStudio Team, 2022). All continuous predictors used in analyses were scaled (means subtracted and divided by the standard deviation). All models investigating duration and frequency parameters included group and bird identity as random effects. Models investigating carol rate and amplitude included only bird identity as a random effect. Group identity was not included as a random effect in these models as it resulted in overfitting and was found to explain minimal variation in the dataset (see Supplementary Materials Table S1, S3). A parallel analysis run with group ID only as a random factor (exlcuding bird ID) confirms that top models do not change.

Only behavioural focals with more than thirty seconds in each noise condition (loud anthropogenic noise/background noise) and with at least one carol during the focal were used to investigate carol rates (*N* = 42 focals from 26 birds across 12 groups). The analysis of factors affecting carol rate used negative binomial GLMMs (with a log link function) to account for overdispersion of data. Predictors included anthropogenic noise condition (loud/background), adult group size and weather condition (clear/cloudy).

For the analysis of factors affecting carol amplitude, we used normally distributed LMMs with anthropogenic noise condition (loud/background), weather condition (clear/cloudy), temperature, humidity, background noise that occurred during the carol (in dB), adult group size, and the number of birds present within 15 meters as predictors.

For the analysis of factors affecting carol duration and frequency parameters, we used normally distributed LMMs with the presence/absence of discernable anthropogenic noise, type of anthropogenic noise (machinery, train, plane, traffic, or no noise), adult group size, and weather condition (clear/cloudy) as predictors.

### Model selection

We conducted model selection using Akaike information criterion values corrected for small sample size (AICc) to determine which candidate models (based on our *a priori* hypotheses) best predicted data patterns (Tredennick et al., 2021). AICc values for each model were recorded and compared. Models were compared to a null model containing only the intercept and random terms. A top model set was constructed containing all models with AICc values within 2 AICc of the model with the lowest AICc, which was considered the most parsimonious model. Terms in the top models were considered significant if their parameter confidence intervals did not intersect zero (Grueber et al., 2011; Symonds & Moussalli, 2011). Where two models had similar AICc values, the simpler model, with the fewest terms contributing to the AICc value, was considered the most parsimonious (Harrison et al., 2018).

## Results

### Carol rate

Individuals carolled significantly less when loud anthropogenic noise was present (5.1 carols per hour), compared to when loud anthropogenic noise was absent (9 carols per hour) (Table 1, Figure 1).

**Table 1.**
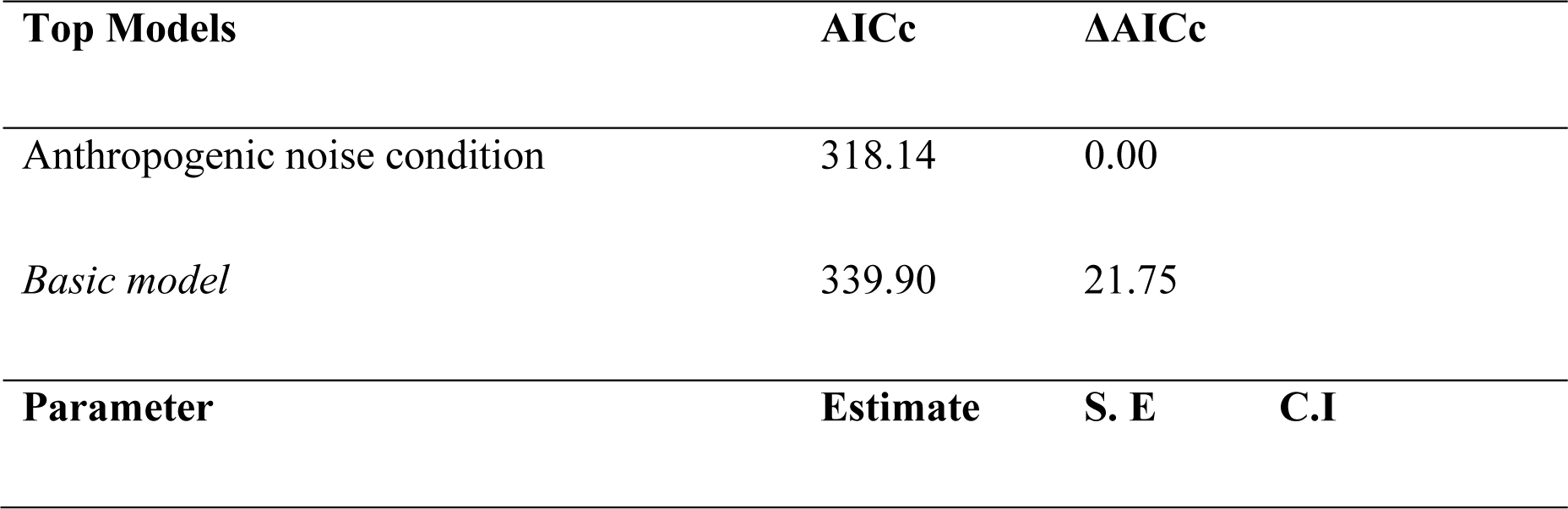
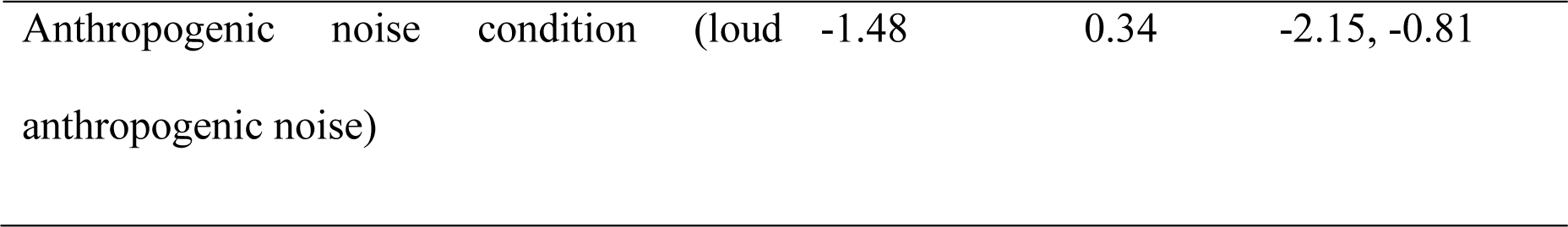
Top model set of candidate terms affecting carol rate (number of carols per 20 minute focal). All models included bird ID as a random term. The top model set includes all models within 2 AICc of the best model and the basic model (intercept-only model) for comparison. Coefficient estimates ± S.E. and 95% confidence intervals (CI) are given below the top model set. *N =* 42 focals from 26 birds from 12 groups. For a list of all models tested, see Supplementary material (Table S2).

**Figure 1.**
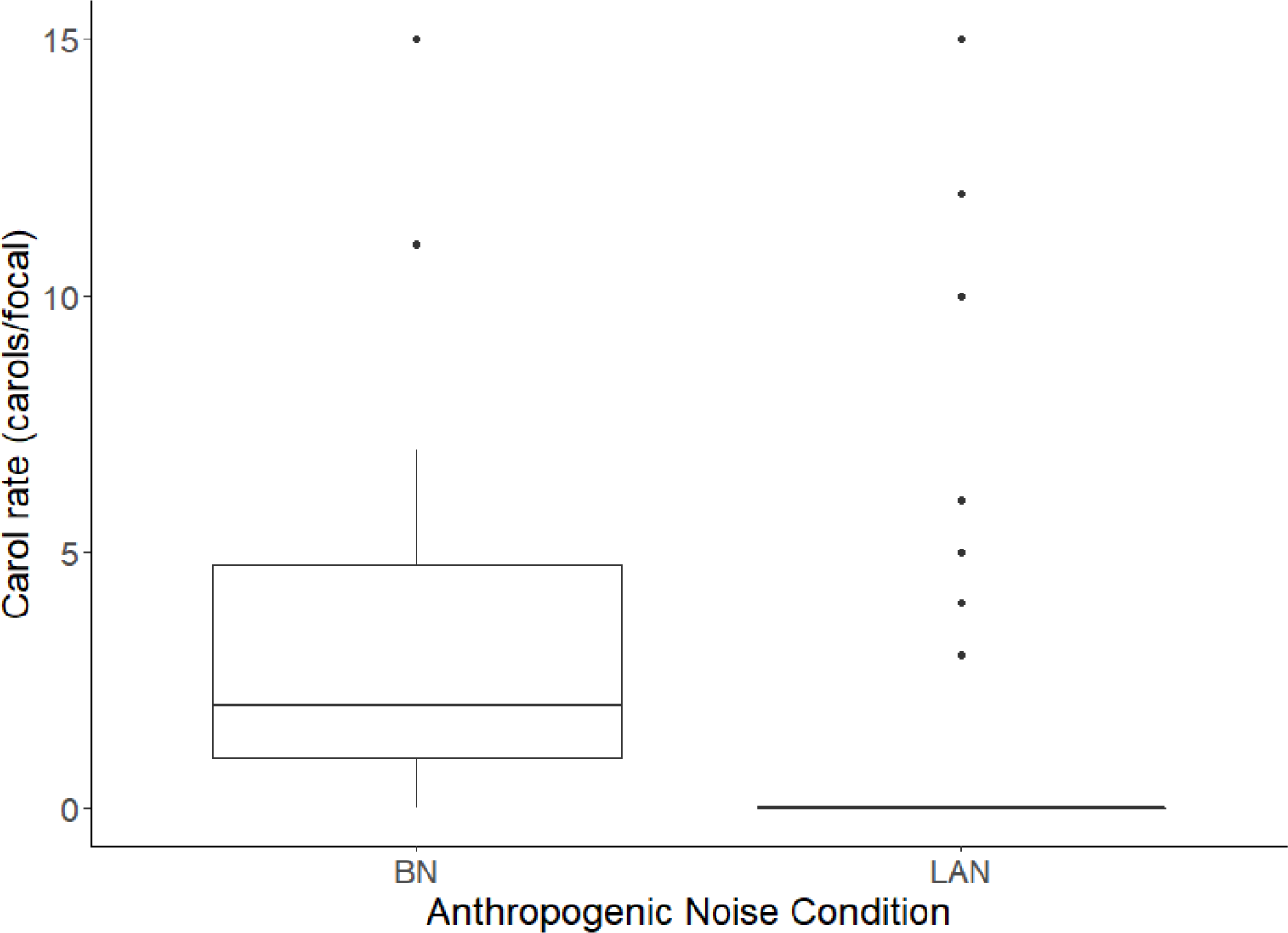
Boxplots show median and quartiles for the distribution of carol rate (number of carols produced per 20-minute focal) in each noise condition (LAN; loud anthropogenic noise, BN; background noise). Outlying points (those > (Q3 + 1.5×IQR)) are represented as circles. *N =* 42 focals of 26 birds from 12 groups.

### Carol amplitude

Carol amplitude was not affected by anthropogenic noise condition (LAN/BN) (Table S4). Carol amplitude was significantly affected by adult group size (Table 2, Figure 2), with birds from larger groups carolling at lower amplitudes compared to birds from smaller groups.

**Table 2.** Top model set of candidate terms affecting carol amplitude. All models included bird ID as a random term. The top model set includes all models within 2 AICc of the best model and the basic model (intercept-only model) for comparison. Coefficient estimates ± S.E. and 95% confidence intervals (CI) are given below the top model set. *N =* 71 carols of 17 birds from 10 groups. For a list of all models tested, see Supplementary material (Table S4).

| Top Models | AICc | ΔAICc |
| --- | --- | --- |
| Adult group size | 411.23 | 0 |
| *Basic model* | 420.05 | 8.82 |

| Parameter | Estimate | S. E | C.I |
| --- | --- | --- | --- |
| Adult group size | -3.09 | 0.64 | -4.34, -1.85 |

**Figure 2.**
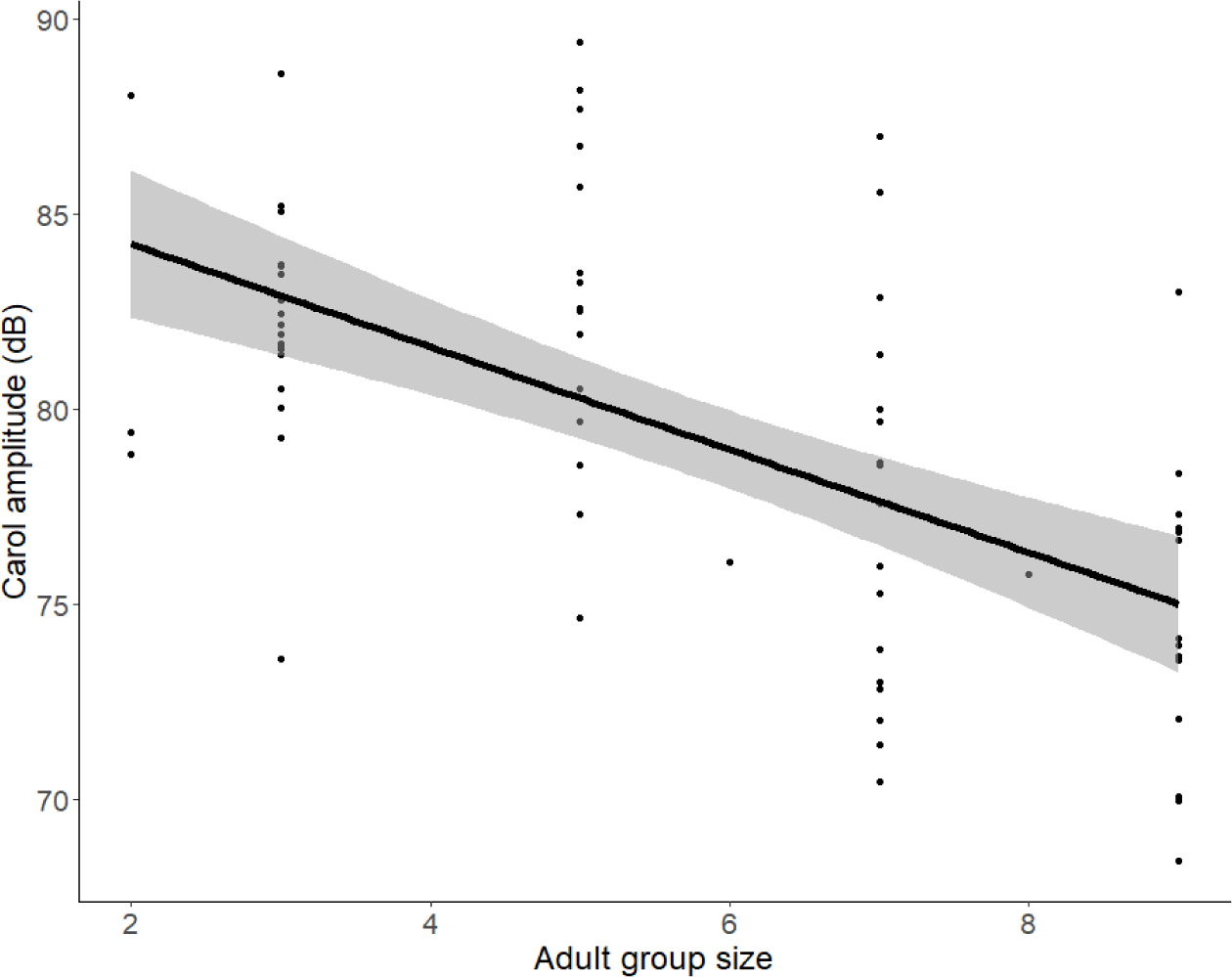
Carol amplitude in relation to adult group size (N = 71 carols of 17 birds from 10 groups). Points represent raw data, fitted line and confidence intervals are from the output of the model presented in Table 2.

### Carol duration and frequency parameters

Carol duration was not affected by the presence of anthropogenic noise, or any other predictor tested (Table S5). However, the presence of anthropogenic noise events significantly affected the peak frequency of carols (Table 4). Carols sung when discernible anthropogenic noise events were present had significantly higher peak frequencies (mean ± SE = 1212.11 ± 42.44Hz) compared to those sung when anthropogenic noise events were not present (mean ± SE = 1070.71 ± 35.50Hz) (Table 4, Figure 5). Minimum frequency, maximum frequency, and frequency bandwidth were not affected by the presence of anthropogenic noise events, or any other predictor tested (Table S7-S9).

**Table 4.** Top model set of candidate terms affecting terms affecting carol peak frequency. All models included group and bird ID as random terms. The top model set includes all models within 2 AICc of the best model and the basic model (intercept-only model) for comparison. Coefficient estimates ± S.E. and 95% confidence intervals (CI) are given below the top model set. *N =* 64 carols of 15 birds across 11 groups. For a list of all models tested, see Supplementary material (Table S6).

| Top Models | AICc | ΔAICc | |
| --- | --- | --- | --- |
| Anthropogenic noise (present/absent) | 174.67 | 0.00 |  |
| *Basic model* | 180.17 | 5.49 |  |
| Parameter | Estimate | S. E | C.I |
| Anthropogenic noise (present/absent) | 0.57 | 0.20 | 0.19, 0.95 |

**Figure 5.**
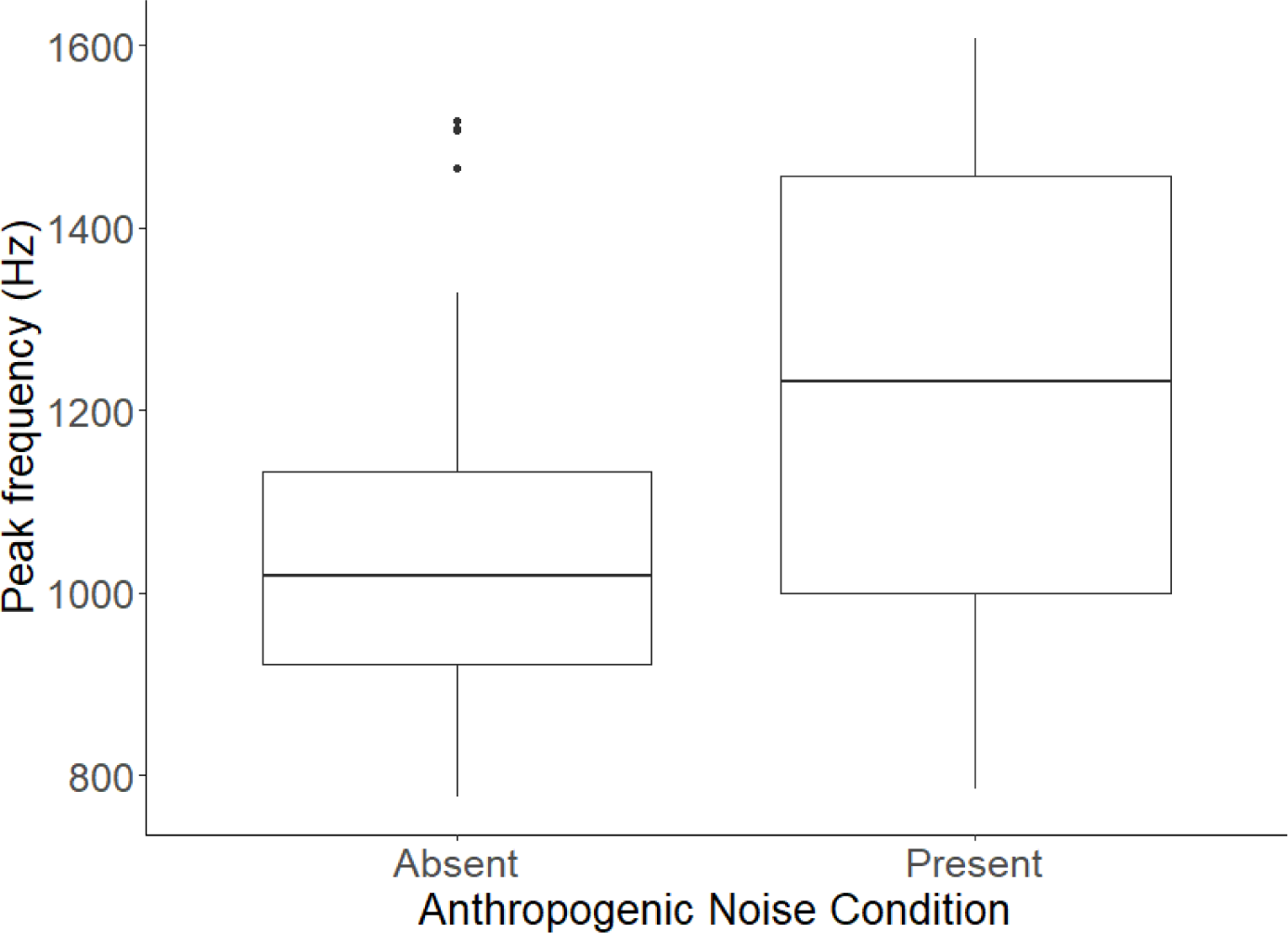
Boxplots show median and quartiles for the distribution of carol peak frequency when anthropogenic noise was present or absent. Outlying points (those > (Q3 + 1.5×IQR)) are represented as circles. *N =* 64 carols of 15 birds across 11 groups.

## Discussion

Our study reveals that anthropogenic noise affects the acoustic structure of female Western Australian magpie carols. When anthropogenic noise was present, magpies reduced their carolling rate, and increased the peak frequency of their carols, but did not alter the minimum frequency of their carols. This finding is surprising, given that recent meta-analyses have suggested it is the minimum frequency of vocalisations that are most often altered by animals when experiencing anthropogenic noise (Duquette et al., 2021; Kunc & Schmidt, 2021). Our results therefore highlight the importance of considering multiple vocalisation parameters, including vocalisation rate, when investigating the effects of anthropogenic noise on acoustic communication, as animal responses to anthropogenic noise are likely to be species specific.

### Carol rate

Individuals significantly reduced their carol rate when loud anthropogenic noise was present. This is consistent with previous work on this species that measured all vocalisations (including alarms, carols, warbles, and mimicry) of individuals with and without noise, and found that individuals vocalised less during loud anthropogenic noise (Blackburn et al., 2023). Such reductions in vocalisation rates during anthropogenic noise have also been found in flying foxes (*Pteropus poliocephalus*) (Pearson & Clarke, 2018), reed buntings (*Emberiza schoeniclus*) (Gross et al., 2010), and several anuran species (Caorsi et al., 2017; Cunnington & Fahrig, 2010), and are hypothesized to occur as a means for individuals to avoid the energetic costs associated with vocalising when their vocalisations are likely to go unheard (Franz & Goller, 2003; Hart et al., 2015). While declines in carol rate during anthropogenic noise may be beneficial for magpies in the short-term, if increases in anthropogenic noise lead to more sustained reductions in vocalisation rates, this could affect social structure and territory defence in this species, as carols play an important role in these aspects of social life (Blackburn et al., 2022; Dutour & Ridley, 2020).

### Carol amplitude

Contrary to our prediction, magpies did not increase the amplitude of their carols when loud anthropogenic noise was present. The absence of the Lombard effect in this species may be because magpies are limited in their ability to increase the amplitude of their carols beyond the 80-90dB threshold recorded in this study (Jakobsen et al., 2021; Zollinger & Brumm, 2015). While the absence of the Lombard effect in magpies may be surprising, a recent meta-analysis concluded this phenomenon was not as uniformly present among species as once thought (Kunc & Schmidt, 2021). Species such as magpies that do not exhibit the Lombard effect may instead utilise other, less energetically expensive means of maintaining communication in noise, such as alterations to the frequency, duration, or rate of their vocalisations, or even non-vocal communication.

Carol amplitude was significantly affected by adult group size, with birds from smaller groups carolling louder than birds from larger groups. Understanding the reason behind the increased carol amplitude of individuals from smaller groups requires an understanding of the function of high amplitude songs in this species. While in some species louder songs are thought to indicate a greater threat level (Brumm & Ritschard, 2011; Ritschard et al., 2012), in others it is instead low amplitude ‘soft songs’ that are posed to convey aggressive intent (Akçay et al., 2015; Searcy et al., 2006). The meaning behind song amplitude therefore appears to be species specific, and future studies are needed to investigate the response of magpies to carols of different amplitudes to determine the function of high amplitude carols in this species, and to understand why females from smaller groups may carol at a higher amplitude.

### Carol frequency parameters

The peak frequency, but not the minimum or maximum frequency, nor the frequency bandwidth of female carols was significantly higher when discernible anthropogenic noise events were occurring. Contrary to our findings, two recent meta-analyses have found that terrestrial animals experiencing anthropogenic noise will generally increase their minimum frequency of their calls (Duquette et al., 2021; Kunc & Schmidt, 2021). Increases in minimum frequency are hypothesized to be important in minimising acoustic masking during anthropogenic noise, as anthropogenic noise is often low frequency, and so will likely disproportionately affect low frequency song elements (Duquette et al., 2021; Slabbekoorn & Peet, 2003). Despite this, increases in the peak frequency of vocalisations during noise have been found in some terrestrial species, including several species of birds (LaZerte et al., 2017; Nemeth & Brumm, 2009; Proppe et al., 2011), and frogs (Cunnington & Fahrig, 2010). Peak frequency is likely to be one of the main determinants of the broadcast distance of a vocalisation, as it is these frequencies that bear most of the energy content of songs (Nemeth & Brumm, 2010; Roca et al., 2016). Therefore, increases in peak frequency may significantly assist magpies in maintaining communication distance during anthropogenic noise events.

## Conclusion

Our study found that female Western Australian magpies alter the rate and peak frequency of their territorial song when anthropogenic noise is present, but surprisingly do not alter the amplitude or minimum frequency of their carols. Changes to the peak frequency of carols could affect the reception and interpretation of these songs by conspecifics, as song frequency has previously been shown to alter receiver response in other species (Brumm & Ritschard, 2011; Huet des Aunay et al., 2014). The findings of this study add to our understanding of the effects of anthropogenic noise on the acoustic communication of wildlife and emphasizes the importance of considering how multiple vocal parameters may change when anthropogenic noise is present.

