## Supplementary Materials Table S1 for "Anthropogenic noise affects vocalisation properties of the territorial song of Western Australian magpies"

***SUPPLEMENTARY MATERIAL***

***Carol rate***

**Table S1.** Comparison of models including bird ID and group ID as random terms. Basic (intercept only) and top model for terms affecting carol rate (number of carols per 20 minute focal) are presented. *N =* 42 focals from 26 birds from 12 groups.

| **Model** | **Random term** | **Variance** | **SD** | **Random term** | **Variance** | **SD** |
| --- | --- | --- | --- | --- | --- | --- |
| Basic (intercept only) | Bird ID | 1.352 x 10^-09^ | 3.677 x 10^-05^ | Group ID | 1.907 x 10^-09^ | 4.366 x 10^-05^ |
| Top model (Anthropogenic noise) | Bird ID | 1.938 x 10^-09^ | 4.402 x 10^-05^ | Group ID | 1.721 x 10^-09^ | 4.148 x 10^-05^ |

**Table S2.** Full model selection output for candidate terms affecting carol rate (number of carols per 20 minute focal). All models included bird ID as a random term. Corrected Akaike information criterion (AICc) and ∆AICc are provided for each candidate model. Only models within 2 AICc of the top model, and with predictors whose 95% confidence intervals did not intersect zero were included in the top model set and are highlighted in bold. *N =* 42 focals from 26 birds from 12 groups.

| **Predictor** | **AICc** | **∆AICc** |
| --- | --- | --- |
| **Anthropogenic noise** | **318.14** | **0.00** |
| Basic | 339.90 | 21.75 |
| Adult group size | 341.57 | 23.43 |
| Weather | 342.00 | 23.86 |

***Carol amplitude***

**Table S3.** Comparison of models including bird ID and group ID as random terms. Basic (intercept only) and top model for terms affecting carol amplitude are presented. *N =* 71 songs of 17 birds from 10 groups.

| **Model** | **Random term** | **Variance** | **SD** | **Random term** | **Variance** | **SD** |
| --- | --- | --- | --- | --- | --- | --- |
| Basic (intercept only) | Bird ID | 13.01 | 3.61 | Group ID | 8.09 | 2.84 |
| Top model (Adult group size) | Bird ID | 1.59 | 1.26 | Group ID | 0.73 | 0.86 |

**Table S4.** Full model selection output for candidate terms affecting carol amplitude. All models included bird ID as a random term. Corrected Akaike information criterion (AICc) and ∆AICc are provided for each candidate model. Only models within 2 AICc of the top model, and with predictors whose 95% confidence intervals did not intersect zero were included in the top model set and are highlighted in bold. *N =* 71 songs of 17 birds from 10 groups.

| **Predictor** | **AICc** | **∆AICc** |
| --- | --- | --- |
| **Adult group size** | **411.23** | **0.00** |
| Temperature | 414.70 | 3.47 |
| Weather | 416.69 | 5.46 |
| Humidity | 416.78 | 5.55 |
| Background noise | 418.26 | 7.03 |
| *Basic* | 420.05 | 8.82 |
| Number of birds within 15m | 421.58 | 10.35 |
| Anthropogenic noise (LAN/BN) | 421.61 | 10.38 |

***Carol duration***

**Table S5.** Full model selection output for candidate terms affecting carol duration. All models included group and bird ID as random terms. Corrected Akaike information criterion (AICc) and ∆AICc are provided for each candidate model. Only models within 2 AICc of the top model, and with predictors whose 95% confidence intervals did not intersect zero were included in the top model set and are highlighted in bold. *N =* 64 carols of 15 birds across 11 groups.

| **Predictor** | **AICc** | **∆AICc** |
| --- | --- | --- |
| **Basic** | **81.41** | **0.00** |
| Adult group size^1^ | 82.80 | 1.39 |
| Anthropogenic noise (present/absent)^1^ | 83.06 | 1.65 |
| Weather | 83.65 | 2.24 |
| Type of noise | 87.72 | 6.31 |

^1^Although within 2AICc of the top model, not included in top model set because confidence intervals intersect 0.

***Peak frequency***

**Table S6.** Full model selection output for candidate terms affecting carol peak frequency. All models included group and bird ID as random terms. Corrected Akaike information criterion (AICc) and ∆AICc are provided for each candidate model. Only models within 2 AICc of the top model, and with predictors whose 95% confidence intervals did not intersect zero were included in the top model set and are highlighted in bold. *N =* 64 carols of 15 birds across 11 groups.

| **Predictor** | **AICc** | **∆AICc** |
| --- | --- | --- |
| **Anthropogenic noise (present/absent)** | **174.67** | **0.00** |
| Weather | 179.93 | 5.26 |
| Basic | 180.17 | 5.49 |
| Type of noise | 180.66 | 5.98 |
| Adult group size | 180.90 | 6.23 |

***Minimum frequency***

**Table S7.** Full model selection output for candidate terms affecting carol minimum frequency. All models included group and bird ID as random terms. Corrected Akaike information criterion (AICc) and ∆AICc are provided for each candidate model. Only models within 2 AICc of the top model, and with predictors whose 95% confidence intervals did not intersect zero were included in the top model set and are highlighted in bold. *N =* 64 carols of 15 birds across 11 groups.

| **Predictor** | **AICc** | **∆AICc** |
| --- | --- | --- |
| Type of noise^1^ | 170.70 | 0.00 |
| **Basic** | **171.42** | **0.72** |
| Weather | 172.82 | 2.12 |
| Adult group size | 173.21 | 2.51 |
| Anthropogenic noise (present/absent) | 173.41 | 2.71 |

^1^Although within 2 AICc of the top model, not included in top model set as it is < 2AICc of the null model.

***Maximum frequency***

**Table S8.** Full model selection output for candidate terms affecting carol maximum frequency. All models included group and bird ID as random terms. Corrected Akaike information criterion (AICc) and ∆AICc are provided for each candidate model. Only models within 2 AICc of the top model, and with predictors whose 95% confidence intervals did not intersect zero were included in the top model set and are highlighted in bold. *N =* 64 carols of 15 birds across 11 groups.

| **Predictor** | **AICc** | **∆AICc** |
| --- | --- | --- |
| **Basic** | **175.48** | **0.00** |
| Weather^1^ | 176.06 | 0.58 |
| Anthropogenic noise (present/absent) | 177.65 | 2.17 |
| Adult group size | 177.73 | 2.24 |
| Type of noise | 182.96 | 7.48 |

^1^Although within 2 AICc of the top model, not included in top model set as confidence intervals intercept 0.

***Frequency bandwidth***

**Table S9.** Full model selection output for candidate terms affecting carol frequency bandwidth. All models included group and bird ID as random terms. Corrected Akaike information criterion (AICc) and ∆AICc are provided for each candidate model. Only models within 2 AICc of the top model, and with predictors whose 95% confidence intervals did not intersect zero were included in the top model set and are highlighted in bold. *N =* 64 carols of 15 birds across 11 groups.

| **Predictor** | **AICc** | **∆AICc** |
| --- | --- | --- |
| Weather^1^ | 177.29 | 0 |
| **Basic** | **178.88** | **1.60** |
| Anthropogenic noise (present/absent) | 180.87 | 3.58 |
| Adult group size | 181.23 | 3.94 |
| Type of noise | 181.41 | 4.13 |

^1^Although within 2 AICc of the top model, not included in top model set as it is < 2AICc of the null model.
